# Phenotyping Root Architecture of Soil-Grown Rice: A Robust Protocol Combining Manual Practices with Image-based Analyses

**DOI:** 10.1101/2020.05.13.088369

**Authors:** P. De Bauw, J. A. Ramarolahy, K. Senthilkumar, T. Rakotoson, R. Merckx, E. Smolders, R. Van Houtvinck, E. Vandamme

## Abstract

**Background:** Breeding towards resilient rice varieties is often constrained by the limited data on root system architecture obtained from relevant agricultural environments. Knowledge on the genotypic differences and responses of root architecture to environmental factors is limited due the difficulty of analysing soil-grown rice roots. An improved method using imaging is thus needed, but the existing methods were never proven successful for rice. Here, we aimed to evaluate and improve a higher throughput method of image-based root phenotyping for rice grown under field conditions. Rice root systems from seven experiments were phenotyped based on the “shovelomics” method of root system excavation followed by manual root phenotyping and digital root analysis after root imaging. Analyzed traits were compared between manual and image-based root phenotyping systems using Spearman rank correlations to evaluate whether both methods similarly rank the phenotypes. For each trait, the relative phenotypic variation was calculated. A principal component analysis was then conducted to assess patterns in root architectural variation.

**Results:** Several manually collected and image-based root traits were identified as having a high potential of differentiating among contrasting phenotypes, while other traits are found to be inaccurate and thus unreliable for rice. The image-based traits projected area, root tip thickness, stem diameter, and root system depth successfully replace the manual determination of root characteristics, however attention should be paid to the lower accuracy of the image-based methodology, especially when working with older and larger root systems.

**Conclusions:** The challenges and opportunities of rice root phenotyping in field conditions are discussed for both methods. We therefore propose an integrated protocol adjusted to the complexity of the rice root structure combining image analysis in a water bath and the manual scoring of three traits (i.e. lateral density, secondary branching degree, and nodal root thickness at the root base). The proposed methodology ensures higher throughput and enhanced accuracy during root phenotyping of soil grown rice in fields or pots compared to manual scoring only, it is cheap to develop and operate, it is valid in remote environments, and it enables fast data extraction.

## 1 Background

Roots are highly important for crop yields as they provide essential functions to the plant, including nutrient acquisition, water uptake, and anchoring into the soil ^1–3^. The spatial distribution, arrangement, and appearance of all root parts and types is collectively referred to as root system architecture (RSA). Optimizing RSA by breeding can enhance crop performance and crop resilience ^4–7^, especially in regions prone to droughts or subject to low inputs (i.e. a limited use of fertilizers or irrigation).

In this context, many root phenotyping platforms have been developed as summarized by Paez-Garcia *et al.* (2015) and Kuijken *et al.* (2015) ^8,9^, each having their advantages and limitations. Some non-destructive phenotyping methods (such as 2D or 3D imaging techniques, visible light imaging, laser imaging, X-ray computed tomography and nuclear magnetic resonance imaging) have been tested to evaluate root architectural traits of plant roots grown in soilless transparent media or rhizoboxes ^8,10,11^. Rhizoscope, a high-throughput system existing of Plexiglas-made rhizoboxes filled with very small glass beads, has been employed in rice root phenotyping to study the number of crown roots, root angle, maximum root length, and nodal root diameter ^12^. However, for breeding purposes, genotype selection or root plasticity should preferably be done based on trait expression under field conditions, as root development can be strongly affected by environmental conditions and genotype × environment interactions (G×E). ^13^

Core and monolith root sampling on field trials in combination with detailed lab analysis of root samples has been deployed in rice studies, but also for other cereal crops. However, root modifications induced by sampling combined with the technical difficulty of accessing soil-grown roots make that the analysis of the whole root system architecture remains challenging. Hence, only few platforms are developed for screening RSA in soil conditions. The strong interactions between genotypes and environmental factors (G×E) ^13^ imply that root phenotyping and trait evaluation should be done in an agronomically relevant environment, which means undisturbed soils under field conditions or in large pots. The collection of root phenotypic data from such relevant environments is costly and labor-intensive, and therefore increasingly becoming a limiting factor in plant breeding (Ahmadi *et al.* 2014; Kuijken *et al.* 2015) ^6,8^. Therefore, there is a strong need for (semi-)automated methods of root phenotyping that are (i) applicable at field scale, (ii) relatively cheap to develop and operate, (iii) easily accessible, and (iv) sufficiently fast so large numbers of individual replicates can be measured.

From all available root phenotyping methods, *Digital Imaging of Root Traits* (DIRT) following the ‘shovelomics’ method of excavation is one of the most simple and robust methods which can easily be applied under field conditions, at any stage during plant development (Trachsel *et al.* 2011; Bucksch *et al.* 2014; Das *et al.* 2015) ^14^-^16^. Despite the destructive character of the method, it is among the few affordable techniques that can be used under multiple field conditions, while enabling the analysis of multiple root traits. Interestingly, DIRT has previously been successfully used for several crops such as maize and beans ^17,18^, but not yet for rice as this was shown to be much more challenging due to the particular complexity of the crop’s root system.

Rice (*Oryza* spp.) is among the most important staple crops in the world, often grown under drought-prone and low-input conditions ^19^. For rice production, the target ecosystems that would thus benefit from optimizing the rice RSA for enhanced production under low inputs include both uplands and rainfed lowlands, while optimizing RSA in high-input irrigated lowlands can also improve the fertilizer- and water use efficiency ^20^. However, while there is a large community focusing on rice breeding, efforts to increase drought resilience and nutrient acquisition efficiency are still constrained by the limited availability of root phenotypic data (Ahmadi *et al.* 2014) ^6^. For a root trait to be a possible target trait when breeding for nutrient acquisition efficiency or drought resilience, it is essential that a sufficiently large genotypic variation in this trait exists, and high-throughput phenotyping methods are thus needed to exploit such genotypic variation. Additionally, phenotyping methods should be able to evaluate the root trait plasticity in response to certain environmental factors. As rice has a highly complex and dense root system, phenotyping of soil grown-rice root systems was shown to be challenging. Therefore, it is necessary to test, validate, and improve the existing semi-automated and high-throughput root phenotyping platforms for rice specifically, in order to enhance the collection of architectural root data from soil-grown rice.

This work aims to evaluate both manual root phenotyping and (semi-)automated image-based root phenotyping (i.e. DIRT) for their ability to differentiate contrasting root phenotypes of rice at field scale. We discuss the challenges and opportunities of imaging soil-grown rice roots, and we propose practical modifications to the method to improve and facilitate future phenotyping of soil-grown rice roots.

## 2 Materials and Methods

### 2.1 Evaluation of rice root imaging under semi-field conditions (large pots)

#### Experimental set-up

Two pot trials (‘pot 1’ & ‘pot 2’) were conducted in a greenhouse at the Sokoine University of Agriculture (6°50’52.8” S, 37°39’31.2”E; Tanzania) in 2016. Details of the experimental set-up and differences among the trials are presented in Table 1 and described in Text S1.a (**Supplementary Information**).

**Table 1:**
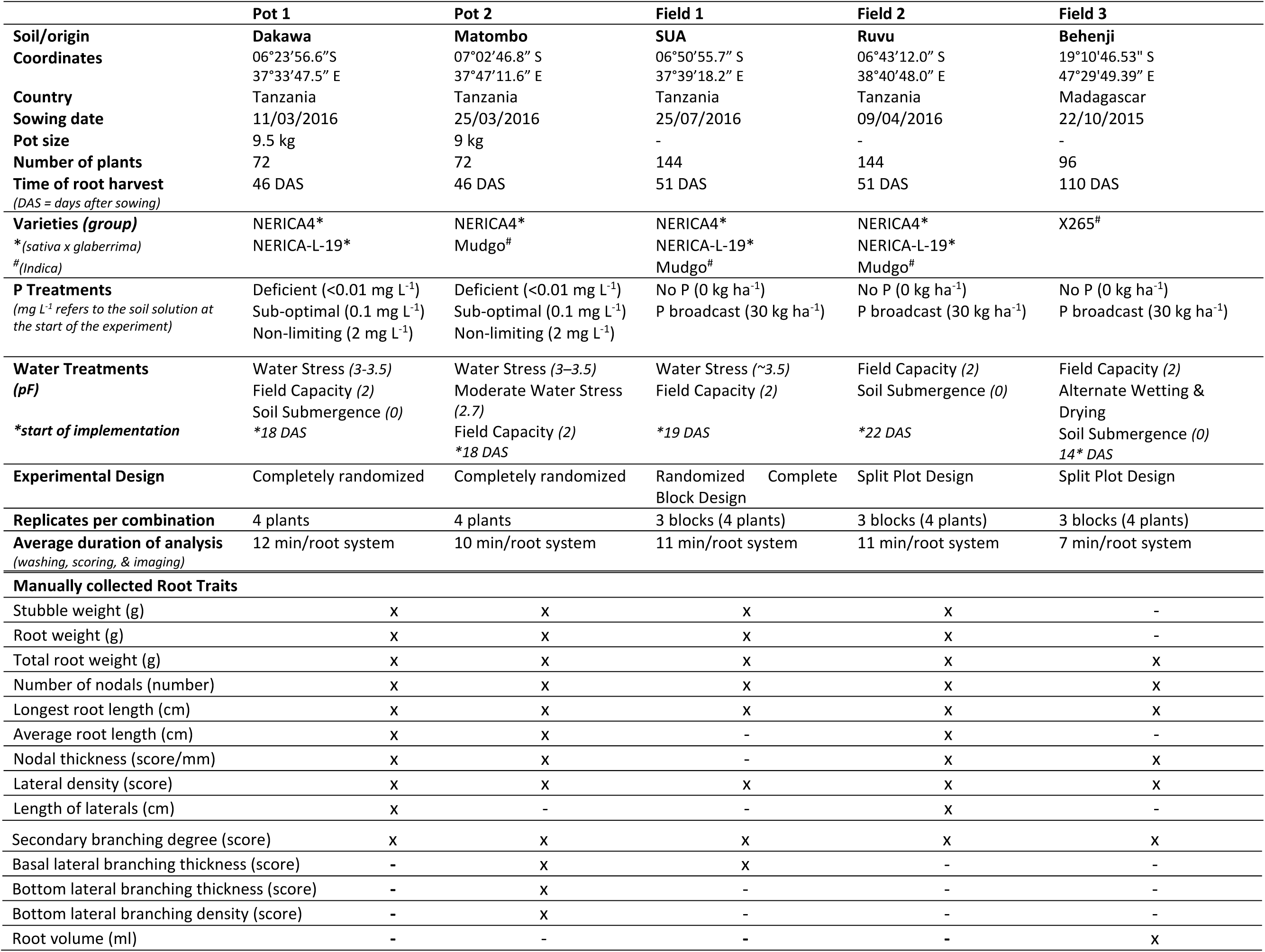
Details and information of the pot and field trials for which root imaging occurred on a dry plate. The manually determined root traits that were collected in each trial are presented with ‘x’ at the bottom.

#### Manual root data collection

At 46 days after sowing (DAS), the root system of each plant was carefully washed out from the soil matrix by soaking and gently shaking the soil matrix in water on a 2 mm net. The root system was then transferred to a dish with clean water where root architectural parameters were manually measured and determined. The number of nodal roots was counted and the average nodal root diameter was measured at the nodal root base, using a transparent ruler (to 0.1 mm). The transparent ruler was placed on the nodal roots, and the average thickness was manually determined. The lateral density, i.e. the spacing of the lateral branches on the nodal roots, was scored using the ‘shovelomics scoreboard’ developed for maize by Trachsel *et al.* (2011) ^15^. The shovelomics scoreboard is a resource for phenotyping roots of soil-grown crops after excavation. For rice, this approach allows the scoring of contrasting densities of S-type laterals by comparing the scoreboard with the density of laterals, as presented on Figure 1 (bottom, right) for the nodal root base (i.e. from stubble up to ca. 15 cm depth). The density was determined by placing the different scoring classes from the scoreboard next to the roots and comparing the densities from the board with the actual density on the root. In some trials these scores of the basal S-type lateral density are complemented with the density of the L-type laterals at the bottom of the root system (below 20 cm depth)). S-type laterals are short and thin lateral roots, emerging at the root base on the nodal roots and they do not have higher order branches. L-type laterals are longer and generally thicker, and they branch further into higher order branches ^21^. These density scores can then be translated into actual values of distance. The secondary branching degree, i.e. the degree of higher order root branching on L-type roots ^21^ evaluated over the whole root system, was manually scored. Lateral root thickness (both at the base and at the deeper roots) were manually scored according to five classes, each corresponding to a thickness class with actual diameter values. The scores used for the different traits are shown in Table S1 (**Supplementary Information**).

**Figure 1:**
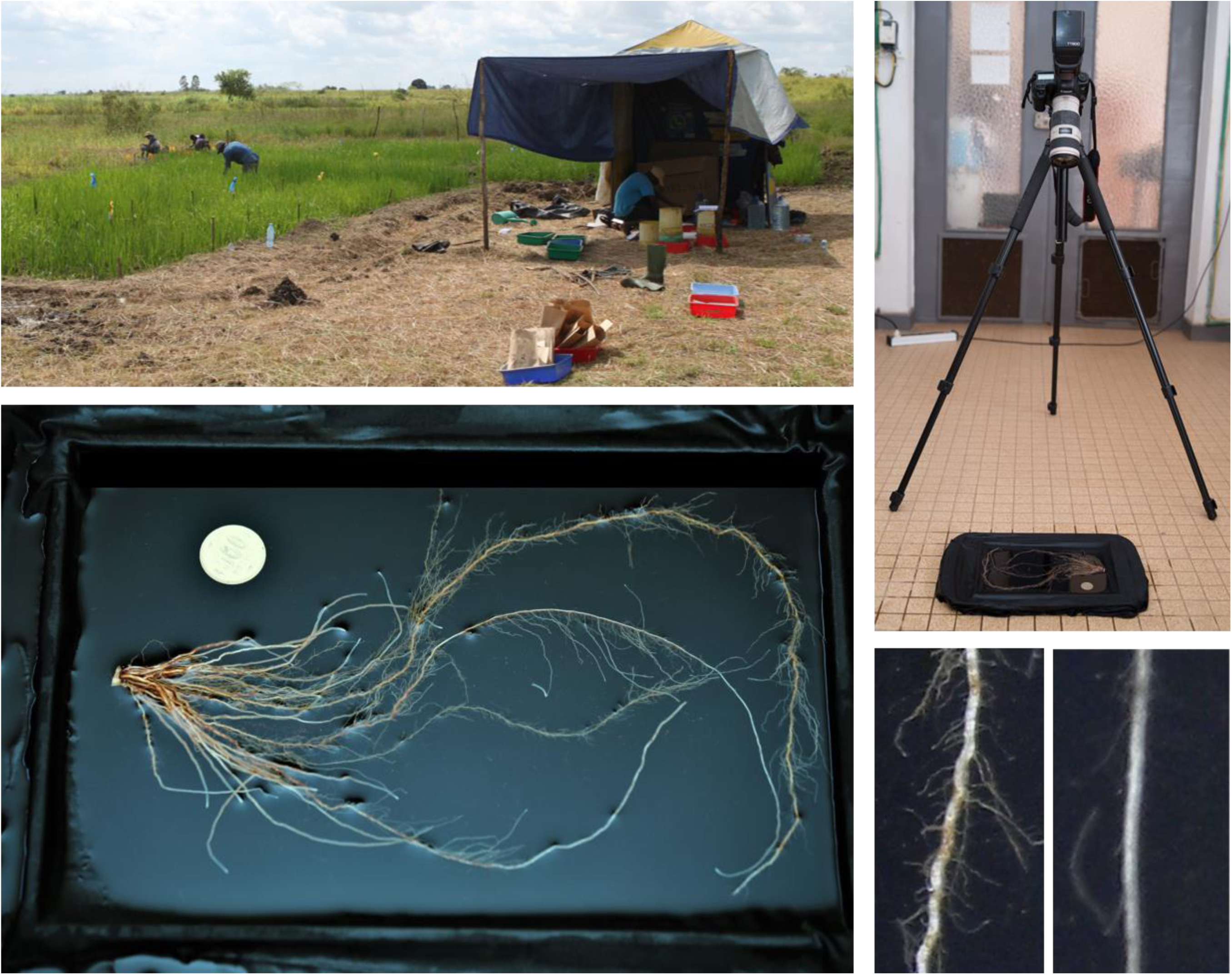
(top, left) Root phenotyping (scoring & imaging) in a temporary tent installed on the field; (top, right) set up of the camera on a tripod to image the roots placed in a water bath; (bottom, left) an excavated rice root system placed in a water bath; (bottom, right) a close up of a nodal root at the root base, displaying a high density score (left) versus a low density score (right).

**Figure 2:**
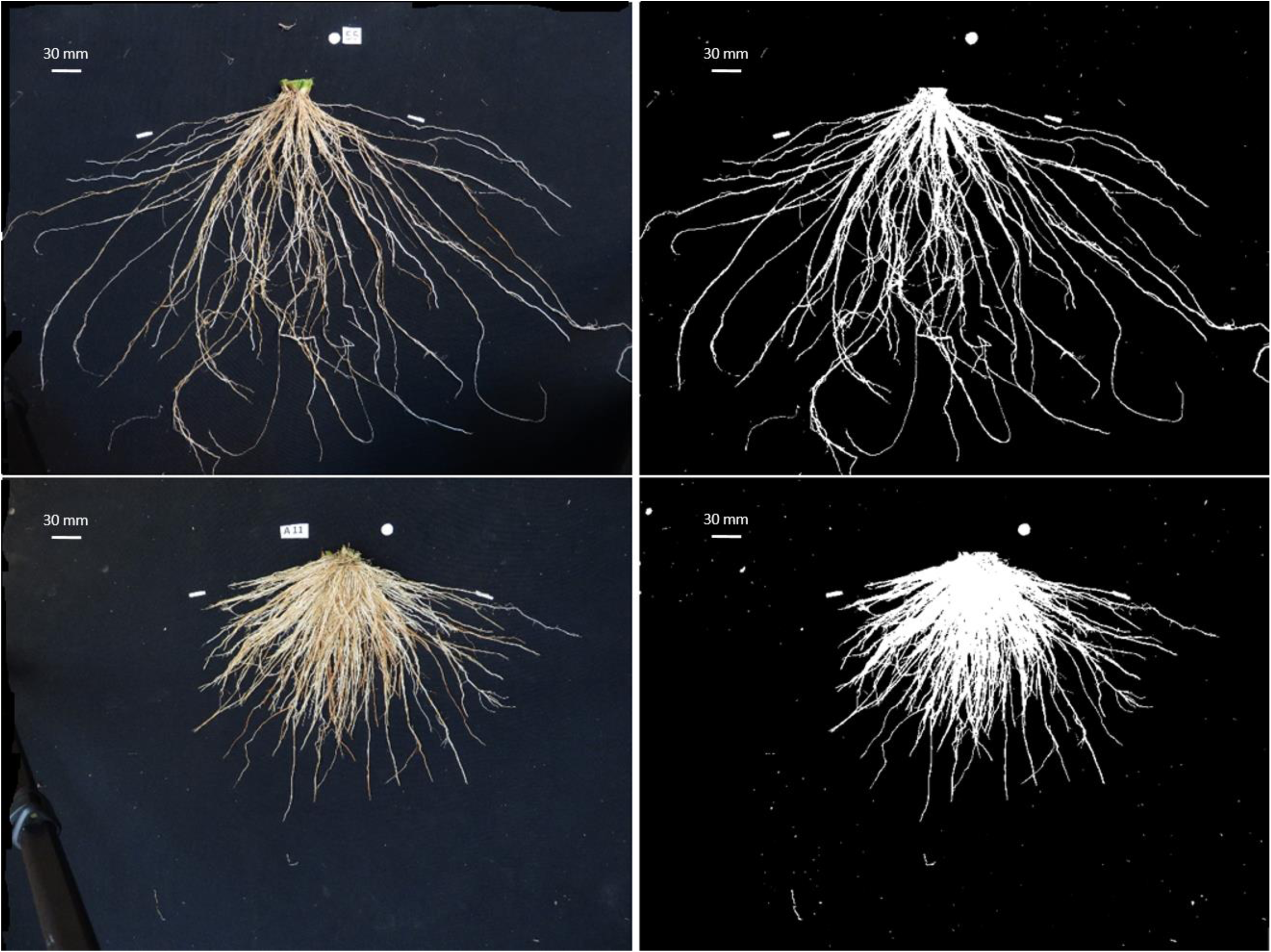
Example of a rice root image (left) and the analyzed black-white picture after conversion by the DIRT software (right). The upper root system was grown in pots (analyzed at 46 DAS), while the lower root system was field grown (analyzed at 51 DAS).

#### Image-based root analysis (DIRT)

After manually determining root architectural traits, the root system was placed on a diffuse black board as described by Bucksch *et al.* (2014) ^16^. A scalemarker with known diameter was provided and images were taken from a fixed distance with a digital camera mounted on a tripod as shown on Figure 1 (top, right). The average duration of root analysis was estimated by dividing the total time needed to analyze all root systems in the trial divided by the total number of analyzed root systems. All images were subsequently processed through the digital image analysis pipeline (DIRT software; http://dirt.cyverse.org/) and all root architectural traits for monocots were extracted. Root traits derived by the imaged-based DIRT software are noted with an ‘*’ in the main text to clarify the origin. After analysis, roots were oven-dried (60°C) and stubble and roots were weighed separately.

### 2.2 Evaluation of rice root imaging under field conditions

#### Experimental set-up and root data collection (manual & image-based)

Two field trials (‘Field 1’ & ‘Field 2’) were conducted in Tanzania while another field trial (‘Field 3’) was conducted in Madagascar (details in Table 1 and Text S1.b (**Supplementary Information**)).

At root harvest (51 DAS for Field 1 & 2 and 110 DAS for Field 3), a soil volume of 20×20×20 cm around each selected rice plant was excavated and roots were washed out from the soil using sieves (2 mm) and water. Root data were collected as described in pervious section, and images of the root systems were similarly taken in a tent temporary placed on the field, as presented on Figure 1 (top, left).

### 2.3 Assessment and validation of manual and image-based root traits

In order to evaluate the capability of the image-based analysis to accurately assess variation in root traits and verify the redundancy of manually determined traits, linear, logarithmic, and exponential relations were first fitted between the image-based traits and the theoretically related root traits that were manually collected (data not shown). Table 2 presents the output variables from the image-based analysis with the corresponding manually determined root traits for which a relation or correlation is theoretically expected based on the nature of the traits. Spearman-Rank correlation coefficients (ρ) between treatment means of the manually determined traits and the related image-based output variables were calculated for each trial, to evaluate whether the image-based analysis was able to accurately differentiate variation and correctly rank performance of root traits among treatments or genotypes, so assessing its potential to replace the manual, more time consuming, determination of a trait. The p-values of these Spearman correlations were computed using algorithm AS 89 for n<1290 and ‘exact = TRUE’ using the cor.test() function in R. These are ‘exact’ for n<10, and use an Edgeworth series approximation for larger sample sizes. ^22^

**Table 2:** Overview of the manually determined architectural root traits (headings) and the closely related image-based traits (extracted from DIRT) below. Based on the nature of these traits, a correlation is expected between the presented traits. The presented output variables from the DIRT software are defined below.

| **Total root weight** | **Number of nodals** | **Nodal root thickness** | **Lateral density** | **Secondary branching degree** | **Longest root length** | **Root volume** |
| --- | --- | --- | --- | --- | --- | --- |
| AREA | RTP_COUNT | TD_MED | RTP_COUNT | RTP_COUNT | SKL_DEPTH | AREA |
|  | AREA | TD_AVG | AVG_DENSITY | AVG_DENSITY |  |  |
|  | SKL_NODES |  | SKL_NODES | SKL_NODES |  |  |
| **DIRT variable** | **Name** | **Definition** |  |  |  |  |
| AREA | **Projected Root Area** | Number of foreground pixels belonging to the root system. Previously defined in GiA Roots. |  |  |  |  |
| RTP_COUNT | **Number of Root Tip Paths** | The overall number of tips detected in the image |  |  |  |  |
| SKL_NODES | **Number of Skeleton Nodes** | The overall number of skeleton nodes in the extracted root shape. (described in Leitner *et al.* 2014) <sup>37</sup> |  |  |  |  |
| TD_MED | **Median Tip Diameter** | Median Tip Diameter |  |  |  |  |
| TD_AVG | **Mean Tip Diameter** | Mean Tip Diameter |  |  |  |  |
| AVG_DENSITY | **Average Root Density** | Ratio of foreground to background pixels within the root shape |  |  |  |  |
| SKL_DEPTH | **Rooting Depth Skeleton** | Rooting depth calculated from the medial axis of the root system. Previously used in GiA Roots. |  |  |  |  |
| DIA_STM | **Stem Diameter** | Stem diameter derived from the medial axis |  |  |  |  |
| SKL_WIDTH | **Skeleton width** | Width calculated from the medial axis of the root system |  |  |  |  |
| ANG_TOP | **Root Top Angle** | Root Top Angle measured between the Random Sample Consensus (RANSAC) fit line at depth of the D10 value and the horizontal soil line. <sup>38</sup> |  |  |  |  |
| ANG_BTM | **Root Bottom Angle** | Root Bottom Angle measured between the RANSAC fit line at depth of the D80 value and the horizontal soil line. |  |  |  |  |
| D10-D90 | **Accumulated width over 10%-90% percent depth (D-values)** | Percentage of width accumulation at 10%-90% depth. The change in width accumulation denotes a change of the root-top angle |  |  |  |  |
| DS10-DS90 | **Slope of the graph of D-values** | Slope of the graph at the D10-D90 values that represents the rate of accumulation |  |  |  |  |

For each root trait (both manually collected and image-based), the ‘*relative phenotypic variation*’ (RPV) was calculated in each trial according to Bucksch *et al.* (2014) ^16^. The RPV of a trait for a given dataset is defined as the ratio between the variance of the trait over all roots in the data set (V_d_) and the average of the trait variances in each specific treatment (*V*_*P*×*W*×*G*_; considering Phosphorus (P), Water (W), and Genotype (G) as experimental factors):

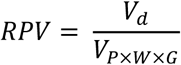

Traits are more likely to be useful in differentiating phenotypic differences when their RPV is larger than 1.

Additionally, for each experiment a multivariate principal component analysis (PCA) was conducted after scaling and centering the meaningful dataset by using the {mixOmics} package in R *version 3.2.2* ^22^. This PCA was conducted for each experiment in order to assess the coherence and relatedness of the multiple contrasting root traits.

### 2.4 An adjusted method to enhance imaging of the rice root architecture

As the rice root system is very dense and sticky (due to the dense branching and very small root diameter of laterals), differentiation among roots and capturing architectural traits by imaging root systems on a dry plate often becomes very challenging. Therefore, we aimed to improve the method of root imaging to enhance the visibility and differentiation of the root types on the image.

#### Experimental set-up and root data collection (manual & image-based)

One pot trial (‘water 1’) was conducted in a greenhouse located at Sokoine University of Agriculture (6°50’53.9”S, 37°39’31.3”E; Tanzania) in 2017, and a second pot trial (‘water 2’) was conducted at the Laboratoire des Radio Isotopes (18°53’56.84”S; 47°33’1.27”E, Madagascar). (Table 3 and Text S1.c (**Supplementary Information**).

**Table 3:**
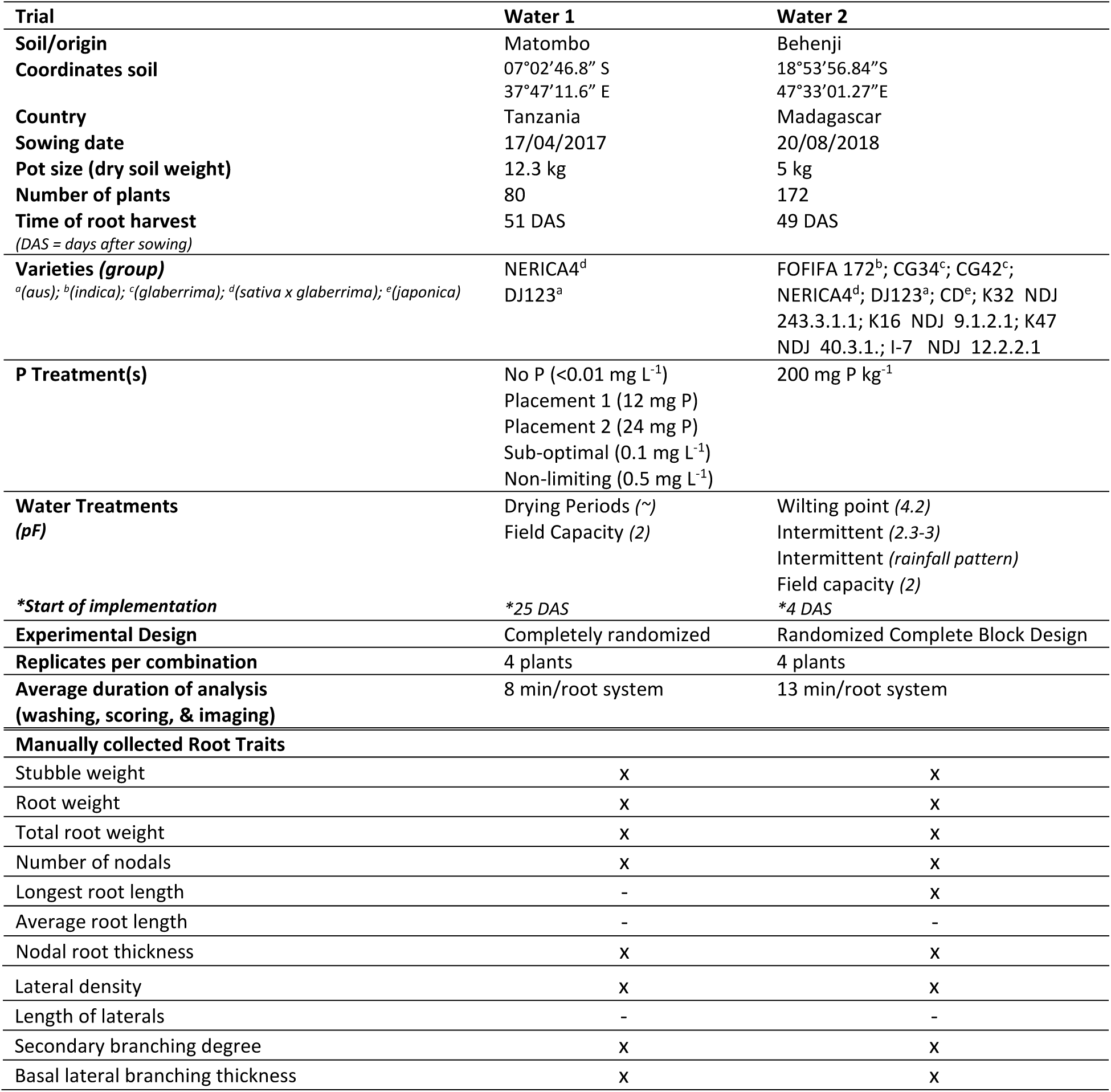
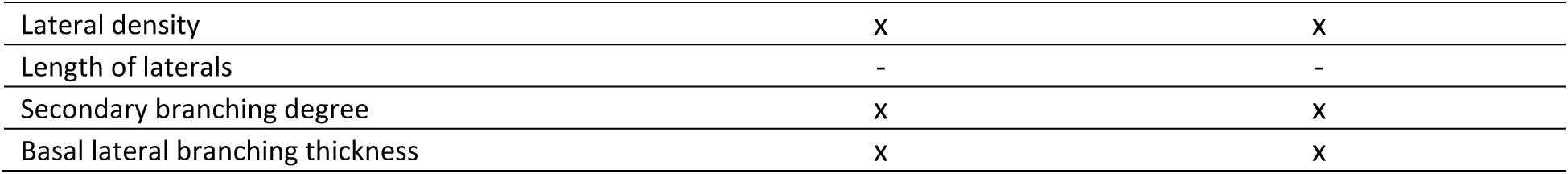
Details and information of the trials for which root images were taken from root systems placed in a water bath. The manually determined root traits that were collected in each trial are presented with ‘x’ at the bottom.

| **Trial** | **Water 1** | **Water 2** |
| --- | --- | --- |
| **Soil/origin** | Matombo | Behenji |
| **Coordinates soil** | 07°02'46.8" S<br>37°47'11.6" E | 18°53'56.84"S<br>47°33'01.27"E |
| **Country** | Tanzania | Madagascar |
| **Sowing date** | 17/04/2017 | 20/08/2018 |
| **Pot size (dry soil weight)** | 12.3 kg | 5 kg |
| **Number of plants** | 80 | 172 |
| **Time of root harvest**<br>(DAS = days after sowing) | 51 DAS | 49 DAS |
| **Varieties (group)**<br><sup>a</sup>(*aus*); <sup>b</sup>(*indica*); <sup>c</sup>(*glaberrima*); <sup>d</sup>(*sativa x glaberrima*); <sup>e</sup>(*japonica*) | NERICA4<sup>d</sup><br>DJ123<sup>a</sup> | FOFIFA 172<sup>b</sup>; CG34<sup>c</sup>; CG42<sup>c</sup>;<br>NERICA4<sup>d</sup>; DJ123<sup>a</sup>; CD<sup>e</sup>; K32 NDJ<br>243.3.1.1; K16 NDJ 9.1.2.1; K47<br>NDJ 40.3.1.; I-7 NDJ 12.2.2.1 |
| **P Treatment(s)** | No P (<0.01 mg L<sup>-1</sup>)<br>Placement 1 (12 mg P)<br>Placement 2 (24 mg P)<br>Sub-optimal (0.1 mg L<sup>-1</sup>)<br>Non-limiting (0.5 mg L<sup>-1</sup>) | 200 mg P kg<sup>-1</sup> |
| **Water Treatments**<br>(pF) | Drying Periods (~)<br>Field Capacity (2) | Wilting point (4.2)<br>Intermittent (2.3-3)<br>Intermittent (*rainfall pattern*)<br>Field capacity (2) |
| **\*Start of implementation** | *25 DAS | *4 DAS |
| **Experimental Design** | Completely randomized | Randomized Complete Block Design |
| **Replicates per combination** | 4 plants | 4 plants |
| **Average duration of analysis**<br>(washing, scoring, & imaging) | 8 min/root system | 13 min/root system |
| **Manually collected Root Traits** |  |  |
| Stubble weight | X | X |
| Root weight | X | X |
| Total root weight | X | X |
| Number of nodals | X | X |
| Longest root length | - | X |
| Average root length | - | - |
| Nodal root thickness | X | X |
| Lateral density | X | X |
| Length of laterals | - | - |
| Secondary branching degree | X | X |
| Basal lateral branching thickness | X | X |

At 51 DAS for ‘water 1’ and 49 DAS for ‘water 2’, the rice root system was washed out and root traits were manually determined as previously described. The root system was then placed in a basin with the same black blanket at the bottom, but filled with water (up to a level of 3 cm). Pictures were then taken from the roots in the water bath from a fixed distance on a tripod, while avoiding reflection on the water surface by placing a light at the lateral side of the basin and positioning the camera in such a way that reflection is avoided (Figure 1 (bottom, left) & Figure 3). The latter can also be facilitated by the use of polarizing filters. Root images were then analyzed by the DIRT software, as previously described for other trials.

**Figure 3:**
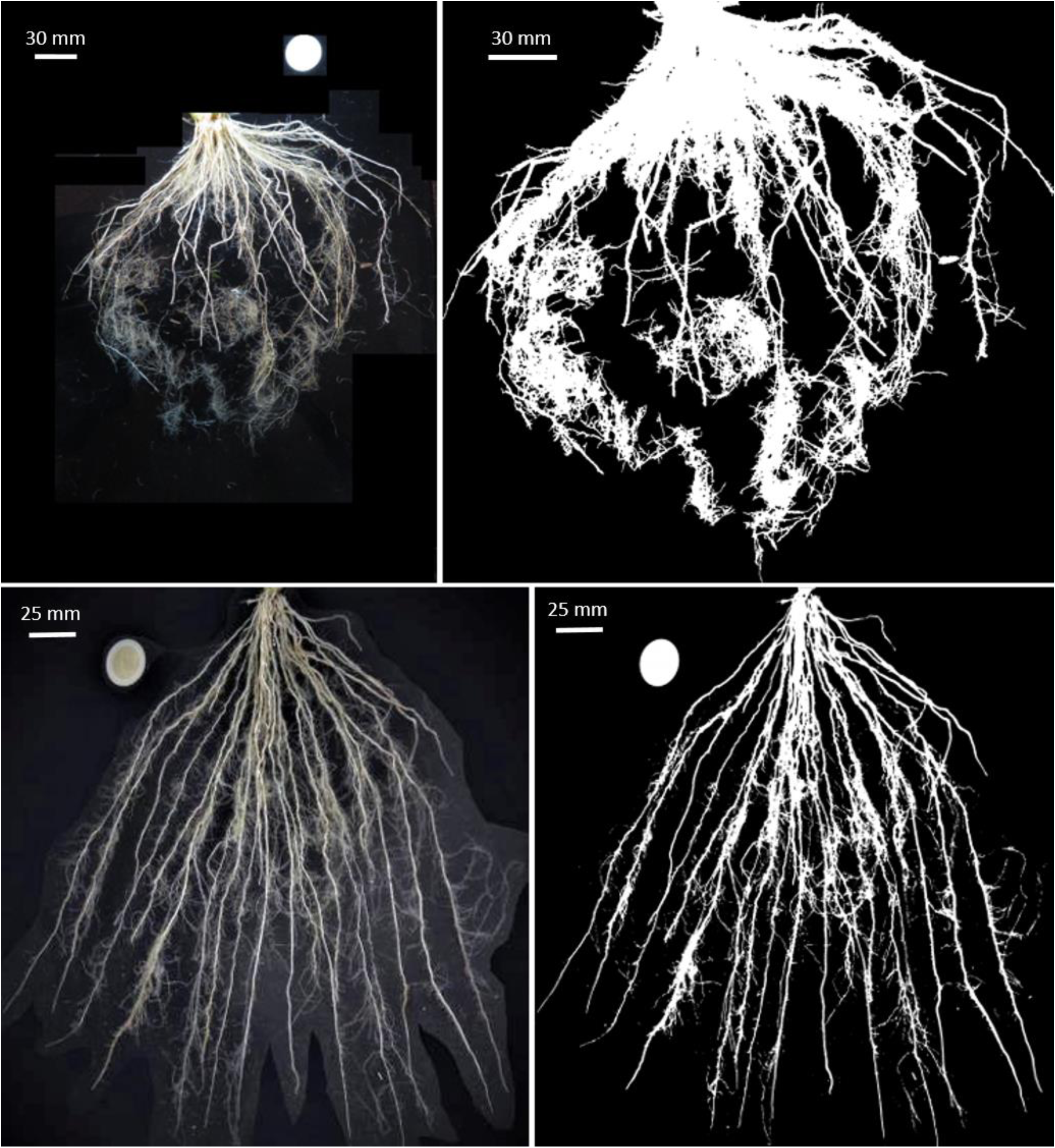
Images of rice root systems taken in a dark water bath before (left) and after (right) conversion by the DIRT software. The upper and lower root system originate from contrasting varieties grown in different trials. Visibility and differentiation of root architecture (laterals and higher order branches) is enhanced by the water bath.

## 3 Results

For each trial, among the image based root traits, strongest relations are demonstrated between the projected root area* and manually determined root traits such as root weight and nodal root number (Table 4). However, the relation between the projected root area* and root weight strongly alters among trials, and the slope tends to increase with increasing plant age but is also larger for field trials compared to pot trials (Figure 4). For all pot trials, an exponential relation is observed between the projected root area* and root weight (Figure 5), while a strong linear relation between nodal root thickness and the mean tip diameter* is observed (Table 4 & Figure 5). The latter relation between nodal root thickness and the mean tip diameter* was not clearly observed in the field trials, possibly following from the fact that many root tips break off during root excavation on the field.

**Table 4:**
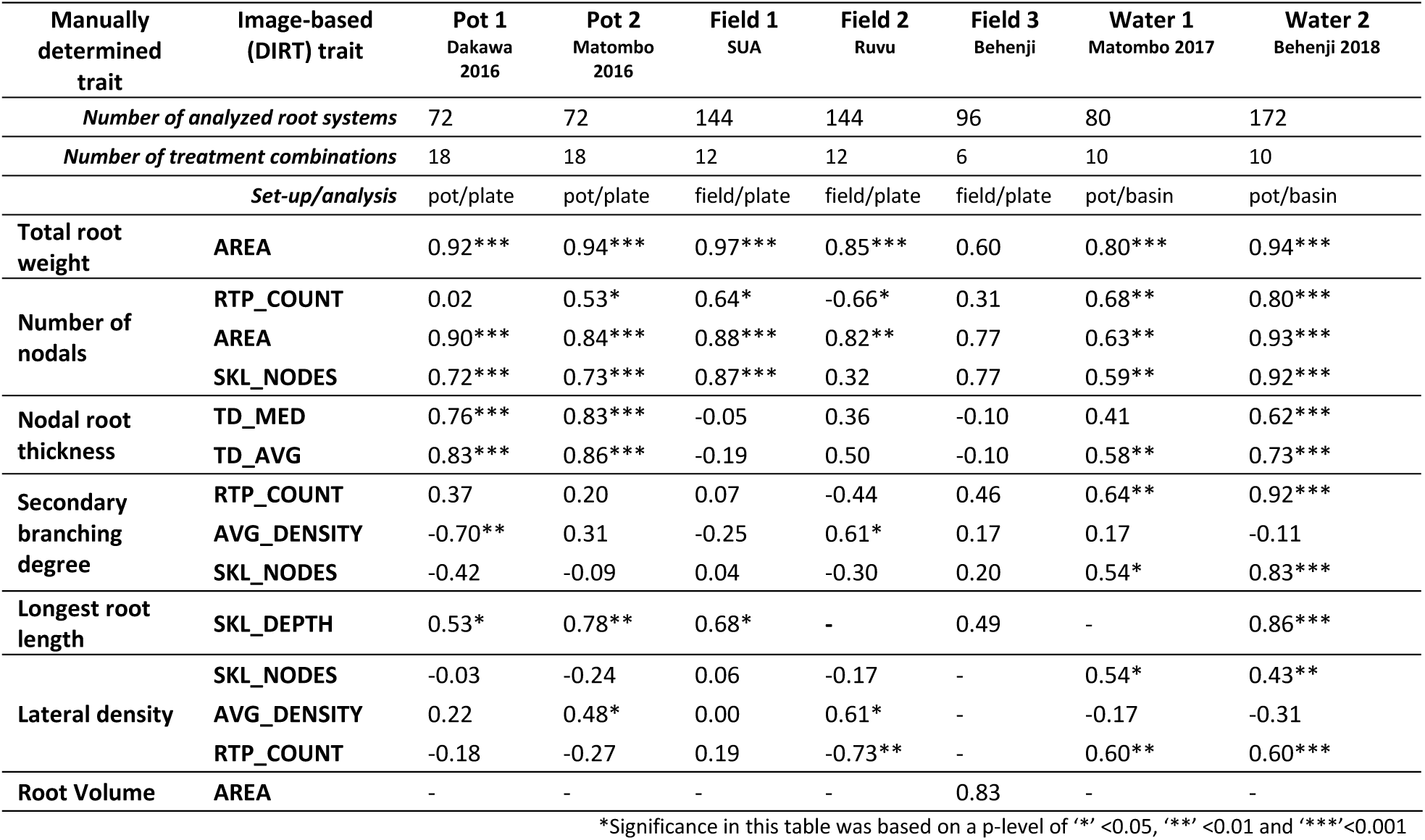
Spearman Rank correlation coefficients (ρ) between the treatment means of manually determined traits and theoretically related image-based output variables (Table 2). The number of analyzed root systems and treatment combinations are also presented. The ‘set-up’ shows whether the experiment was conducted in pots or on the field, while ‘analysis’ refers to the method of root imaging (i.e. on a dry plate or in a water bath)

**Figure 4:**
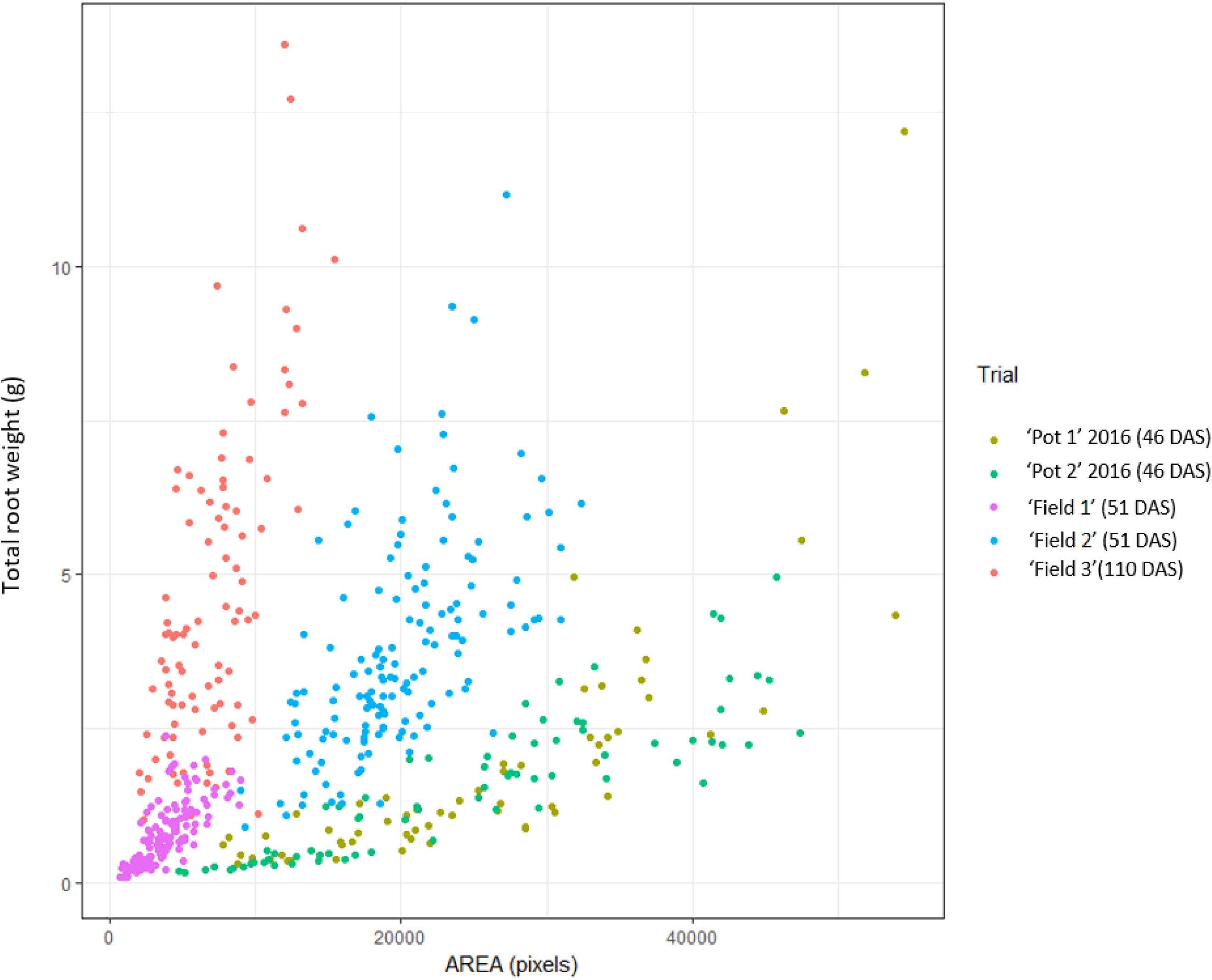
The analyzed AREA (pixels) versus Total root weight (g) for all experiments for which root images were taken on a dry plate with a black diffuse background. The figure illustrates the inconsistent relationships between the two variables among experiments and an increasing slope with plant age.

**Figure 5:**
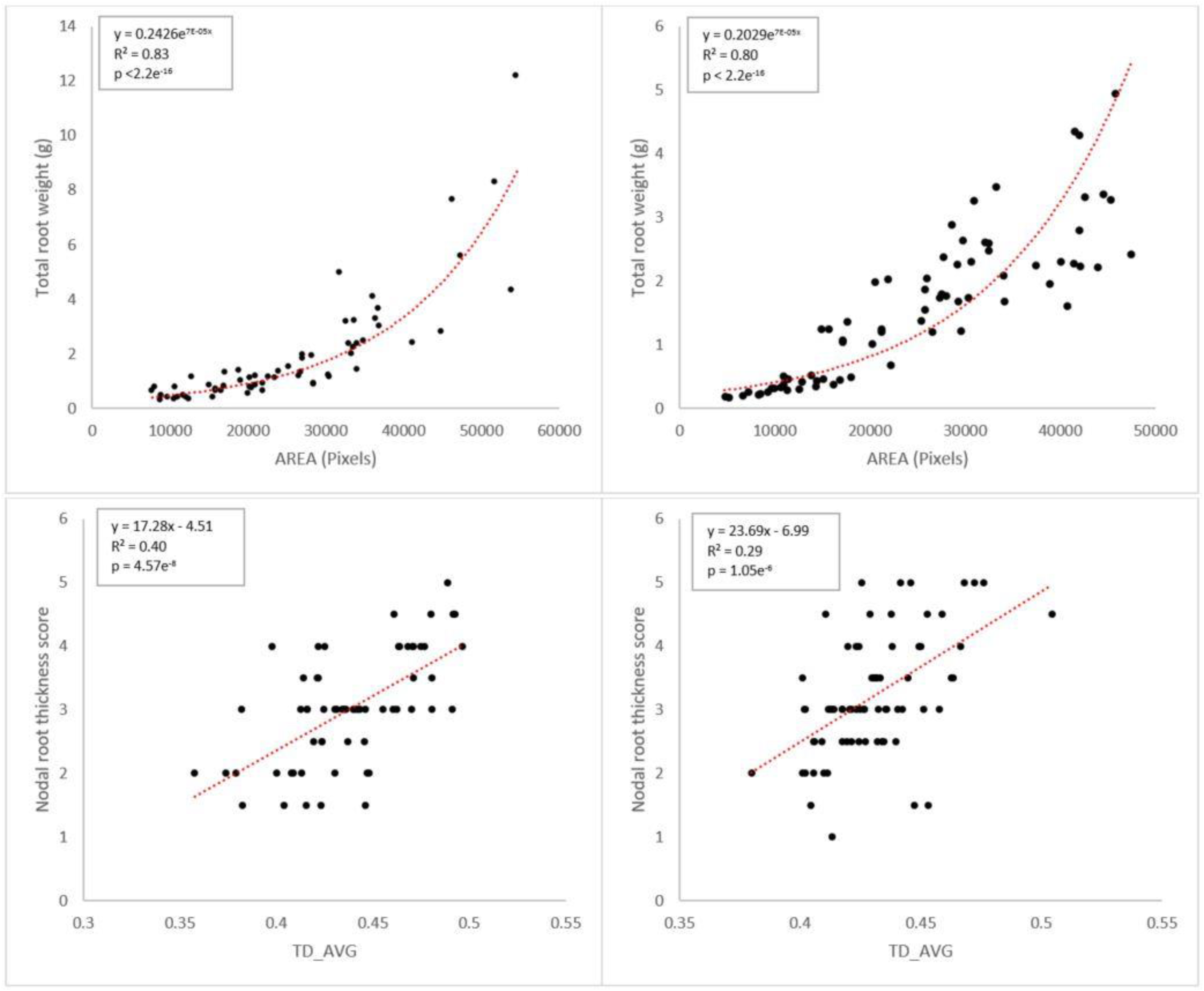
Relations between the projected root area* (pixels; AREA from the DIRT-output) versus Total root weight (g) (at the top), and between the mean tip diameter* (TD_AVG; from the DIRT-output) and the manually determined nodal thickness score for ‘Pot 1’ (Dakawa 2016) (left) and ‘Pot 2’ (Matombo 2016) (right). The relation details, R^2^, and p-values of each fit are presented at the top.

Generally strong correlations are indeed observed between the total root weight and the projected root area*, and between the number of nodal roots and the number of root tip paths*, the number of skeleton nodes*, and the projected area* (Table 4). Interestingly, the longest root length strongly correlates to the rooting depth skeleton* analyzed on the image. Inconsistent correlations among the trials are observed between image analysis derived traits and the manually determined traits ‘secondary branching degree’ and ‘lateral density’. However, root imaging in water strongly improved the correlations of the latter two manually determined traits (i.e. ‘secondary branching degree’ and ‘lateral density’) with the number of skeleton nodes* and the number of root tip paths*, while it did not improve the correlations with the average root density*. Nodal root thickness shows a strong correlation with the mean tip diameter* and the median tip diameter* for all trials, except for ‘Field 1’ and ‘Field 3’ (Table 4).

Interestingly, RPV values of manually determined root traits are generally larger than the image-derived root traits (Table 5). Highest RPV values are observed for the manually determined root traits: total root weight (RPV from 2-6.6), number of nodal roots (RPV from 1.7-12.6), secondary branching degree (RPV from 1.1 – 5.3), lateral density (RPV from 1.2-5.6), and nodal thickness (RPV from 1.1-10.5). Image-based traits with a consistent relatively high RPV are the number of skeleton nodes* (RPV from 1.2-4.8), *AREA* (RPV from 1.4-6.3), and the number of root tip paths* (RPV from 1.1-3.6). Trends in RPV values are less consistent among trials for the stem diameter* (RPV from 1.0-2.7), the average root density* (RPV from 1.0-1.8), the rooting depth skeleton* (RPV from 1.0-2.2), the mean tip diameter* (RPV from 1.0-1.9), and the skeleton width* (RPV from 1.0-5.8). RPV values from the accumulated widths over 10%-90% percent depth* (*D10-90)* and the slopes of the graphs of these D-values* (*DS10-90)* are consistently close to 1. (Table 5 & Figure S1 (**Supplementary Information**))

**Table 5:**
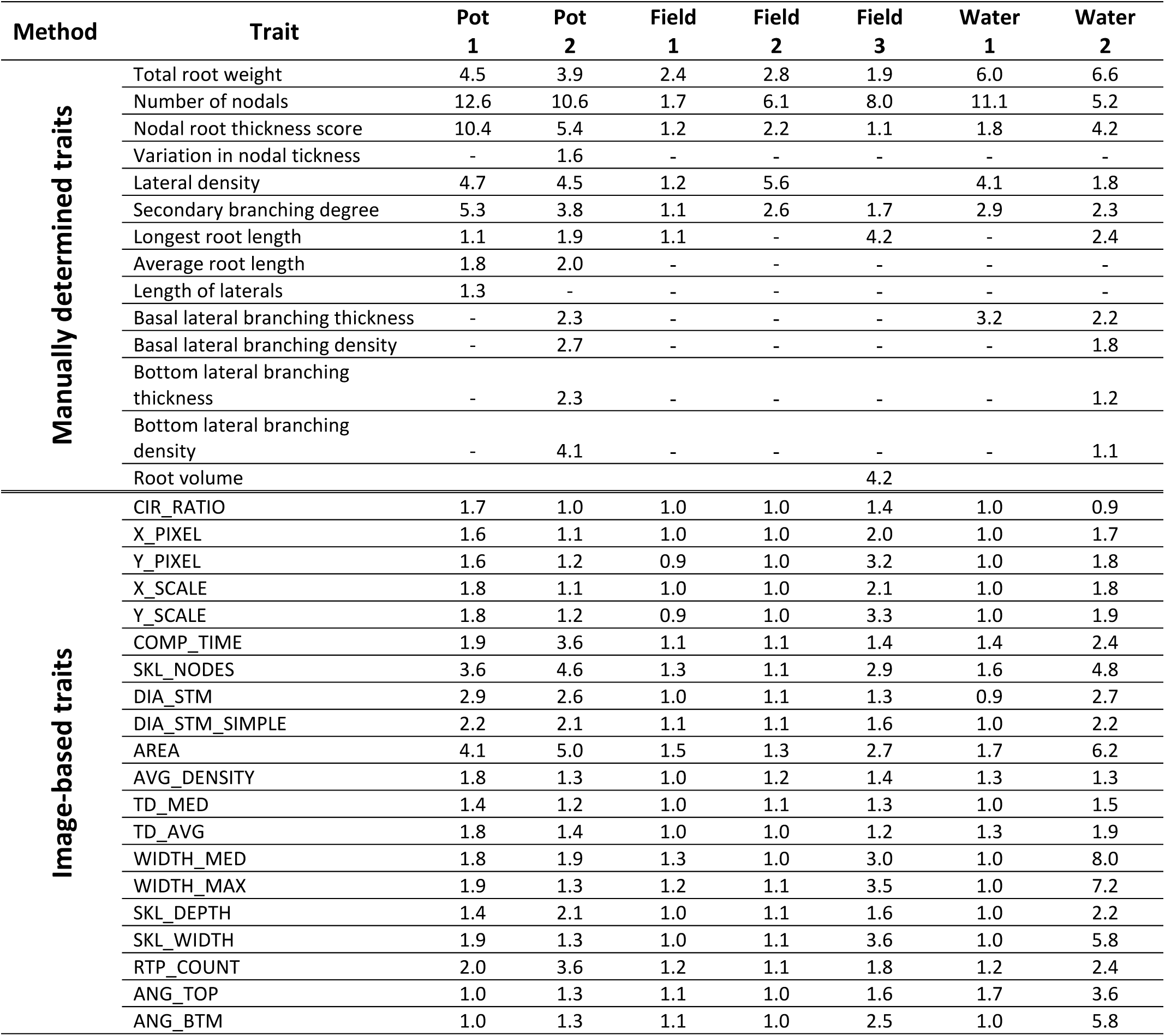
The relative phenotypic variation (RPV) of each root trait determined for each experiment. Traits presented are both manually determined (top) and calculated based on images (bottom) (Das et al. 2015)^14^. Traits are more likely to be useful in differentiating phenotypic differences when their RPV is larger than 1. The explanations of the image-based trait names are presented in Table 2.

Results of the PCA highlight the inherent correlations between several root traits. The first principal component explained 16 to 45% of the variation while the second component comprised 6-20% of the variation in the different experiments (Table S2, **Supplementary Information**). Generally, principal component 1 was predominantly loaded by size-related phenes such as root weight, the projected root area*, and number of nodals, while loadings on PC2 were more variable and mainly presented by secondary branching degree and lateral density (Figure S2, **Supplementary Information**). Figure 6 demonstrate how a PCA of the suggested root traits (combined manual and image-based) enables to distinguish and group root phenotypic performance related to genotype selection (in experiment Water 1) or water treatments (in experiment Water 2), highlighting the power and applicability of this phenotyping protocol.

**Figure 6:**
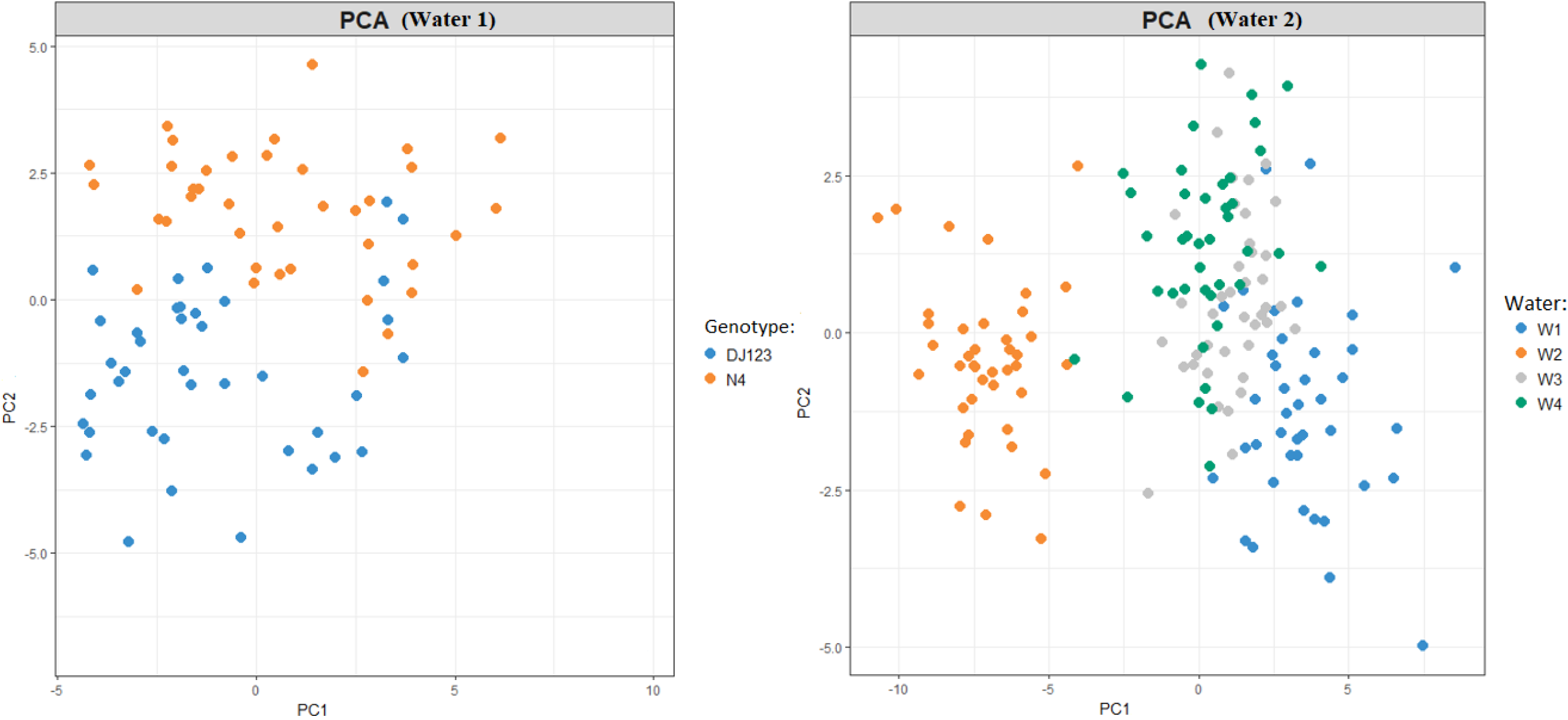
Score plots of the Principal Component Analysis from the root data combining manual scoring and image-based root analysis. These plots indicate how the proposed method presented in this chapter enables to differentiate contrasting root phenotypes, here coinciding with contrasting genotypes in trial ‘water 1’ (Left), and different water treatments in trial ‘water 2’ (right).

## 4 Discussion

### 4.1 Challenges and opportunities for image-based root phenotyping of soil-grown rice

We have evaluated both manual and image-based root phenotyping methodologies for soil-grown rice in multiple environments. The inconsistent relationships between some manually determined and image-based traits across all experiments demonstrate that a reliable and consistent quantification of root traits (e.g. root weight (g), nodal thickness (mm), depth (cm)) based on image-based output variables remains very challenging. Variations induced by environmental characteristics, plant age, and harvesting method should definitely be considered when comparing results from different environments and this variation precludes a robust trait quantification based on images. Variability among root phenotypes (from one genotype) across several environments and experiments was also demonstrated by Trachsel *et al.* (2011) ^15^ for maize, and this can be explained by the strong genotype-by-environment interactions ^12,23^.

The rice root system is very dense and has a complex structure (Rebouillat *et al.* 2009; Ahmadi *et al.* 2014) ^6,24^, and the large degree of root overlap forms a major challenge for image-based root analysis. Figure 4 shows the variable relations between the projected root area* and total root weight, induced by environment and plant age. The steeper slope of the relation between the projected root area* and total root weight for older plants is explained by the increasing overlap of roots at the nodal root base combined with a larger stubble weight in older plants. This increasing root overlap with increasing root size following a larger nodal root number additionally results in exponential relations between the projected root area* (exponent) and total root weight as shown in Figure 5. Additionally, in the field only shallow roots can be collected (seen on Figure 2), and more root overlap occurs at this root base. Hence, this results in a smaller projected root area* relative to the root mass of field-excavated root systems (i.e. steeper slopes for field experiments on Figure 4). Thus, it should be considered when analyzing larger root systems in field conditions that the overlap of nodal roots at the root base can decrease the accuracy of the image based analysis and therefore it would underestimate size related traits.

As rice roots are relatively flexible, the root angle is modified during washing and placing the roots on the board. Therefore, root angle and other ‘angle-related’ DIRT traits as the skeleton width*, the root top angle*, the root bottom angle*, the accumulated widths over 10%-90% percent depth* *(D10-90)*, and the slopes of the graphs of these D-values* *(DS10-90)* are considered as inaccurate and unreliable parameters without biological meaning. These parameters cannot be determined correctly by the shovelomics method of root excavation and hence they were excluded from the analyses. To overcome this challenge of root angle modifications during excavation, the root angle could be analyzed in field conditions by the ‘basket method’ ^25,26^, using vertical core sampling ^27^, or it could be analyzed after washing out a vertical plain along the root ^28^.

An additional related challenge for rice root phenotyping is the ‘stickiness’ of the fine laterals. The architecture of laterals becomes invisible as soon as the root system is taken out of water. Therefore, the extraction of root system architectural traits by ‘*Root Estimator for Shovelomics Traits (REST)*’ as described by Colombi *et al.* (2015) ^17^ or 3D imaging in a box ^29^ would not work successfully for rice.

Despite some inconsistent relations among manually determined and image-based traits, the high Spearman rank correlation coefficients between several traits (Table 4) highlight the possibility to differentiate root phenotypes by comparing treatment means of several image-based traits within a trial. The Spearman rank correlations indicate that some image-based traits can indeed replace the manual collection of certain traits, in order to speed up the phenotyping process on the field, without losing useful information. Interestingly, pooling all analyzed traits into composite descriptors after a principal component analysis allows the identification of distinct root phenotypes, related to treatments or genotypes (Figure 6), highlighting the power of this method.

Among the image-based traits considered in our study, the projected root area* can be used most successfully to rank phenotypes according to root size, and this probably follows from two underlying reasons. First, it is the trait that can most accurately be captured on the image – thus with lowest measurement noise. Second, it can be considered as a size descriptor that indeed integrates several branching and vigor information, being meaningful for identifying distinct root phenotypes. Additionally, the mean tip diameter* and the median tip diameter* can also successfully replace manual measurements of the root tip diameter when root systems are not too old and dense (Figure 5 & Table 4). However, additional care should be taken to the differences between root tip thickness and nodal thickness at the root base, as the latter trait cannot accurately be determined by the image based DIRT analysis. The rooting depth skeleton* can successfully be used to analyze the longest root length of the excavated root system, but attention should be paid to the challenges of excavating deep roots on the field and correctly placing the longest root in the root bath without folding it. Roots of field-grown rice may grow deeper than 1 m ^30^, a depth that often precludes excavation. Deep rooting of rice was therefore suggested to be analyzed in pots ^31^, or by soil coring and raised field beds ^25^. The number of skeleton nodes* and the number of root tip paths* also showed potential to differentiate phenotypes, and when imaging rice roots in water the correlation with the number of nodal roots, lateral density, and secondary branching degree increased. The number of skeleton nodes* and the number of root tip paths* integrate the influences of both the nodal root number, lateral density, and secondary branching degree, but the image-based method is not able to identify or distinguish these particular traits. The latter three manually determined traits can thus not be replaced by one single image based trait. This shows that the image-based analysis is not accurate enough to capture all details required for differentiating genotypes at a more subtle scale of architectural traits. This does not invalidate the DIRT approach, but it simply shows its limitations. Therefore, it would be beneficial to combine both manual and image-based methodologies when phenotyping rice roots, as proposed in the next section.

The large values of the relative phenotypic variation (RPV) for some manually determined and image-based traits highlight the power of a combined method to differentiate contrasting root phenotypes related to genotypes or environmental factors. Interestingly, the manually determined traits show generally larger RPV values than the DIRT traits, which is probably related to the reduced measurement accuracy of image-based traits, mainly explained by the strong root overlap on images. Interestingly, the lateral density and secondary branching degree showed a consistently high RPV value in most trials, indicating large variations of this trait among genotypes or treatments^12,13^, and this cannot clearly be detected by the image-based analysis. Basal S-type laterals cannot be differentiated on the images because they are very thin and hidden by overlapping nodal roots at the base. Hence, as this trait displayed a high differentiation potential among genotypes or environmental factors, we argue that this basal lateral density should be manually determined on the root system or by analyzing images of single excised roots. Additionally, previous studies have previously shown that such manual analyses of lateral density and secondary branching can indeed successfully assess phenotypic variation in rice root performance ^32,33^.

Interestingly, the PCA demonstrated that most variation in all experiments is explained by size-related phenes, mainly dominated by the image-based trait, projected root area*. Such high variance induced by size related root phenes was also observed for maize by York and Lynch (2015) ^34^ and it indicates that there is indeed an underlying genetic basis causing correlations among all size related traits. This implies a pleiotropic genetic control of several size related root phenes, such as a general root vigor driver for PC1 ^34^. Interestingly, the manually determined traits basal lateral density and secondary branching degree were found to be independent from size related traits and this observation indeed suggests a different genetic program of lateral branching (i.e. lateral primordia initiation and their subsequent emergence), independent from root vigor ^35,36^.

### 4.2 Future prospects and recommended protocol for rice root phenotyping

We have shown that image-based analysis of the rice root system architecture can be successful for several traits, but not all traits are reliable and useful. Concerning the previously discussed challenges and opportunities for rice root phenotyping in field conditions, we now propose a phenotyping method that combines both the manually determination of few root traits with an adjusted method of root imaging, as presented in Figure 7.

**Figure 7:**
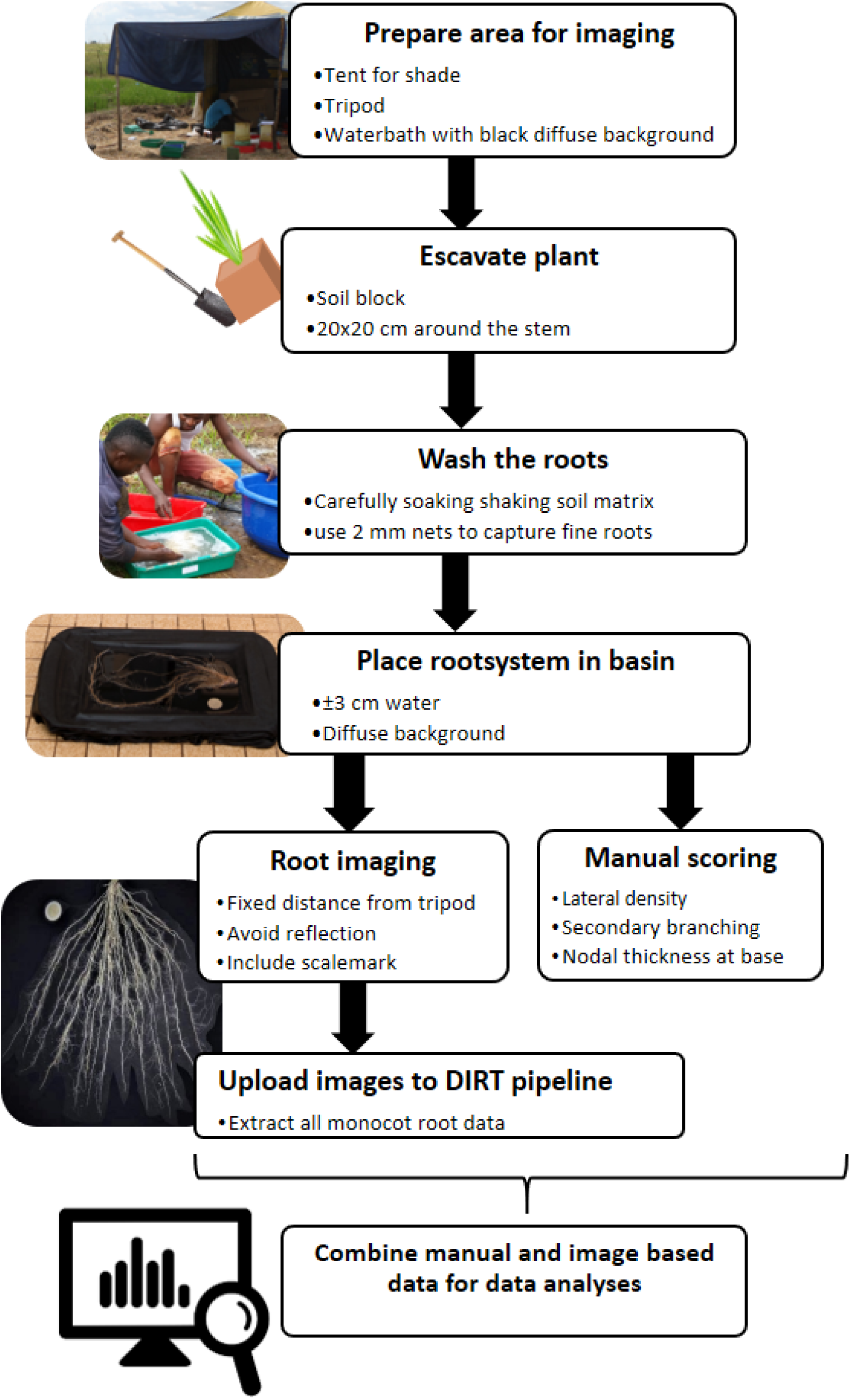
A schedule presenting the suggested phenotyping protocol for rice, applicable at field scale. It combines root imaging in water with some manual analysis.

We suggest to manually determine the lateral density, secondary branching degree, and the nodal root thickness at the root base (if relevant for the study) before imaging the root system. In order to increase resolution of the image based analysis and enhance differentiation among nodal and lateral roots, we then suggest to take root images in a water bath (ca. 2-3 cm depth) on a diffuse black background, before analyzing them with the DIRT software. Root traits as the projected root area*, the mean tip diameter*, the rooting depth skeleton*, the average root density*, the number of skeleton nodes* and the number of root tip paths* can then be successfully extracted. Combining these three manually determined traits with the image-based analysis from root images in a water bath would efficiently and most accurately cover the widest range of interesting root traits, and this method has indeed shown to have a high potential of differentiating root phenes (Figure 6).

Attention should be paid to the strong root overlap when roots are becoming older, larger, and very dense. Future research would therefore benefit from identifying the earliest phenological stage at which rice plants show root architectural traits also expressed at later growth stages and being independent from size (as discussed before). Harvesting younger plants would facilitate the differentiation of root architectural traits by imaging, reduce the efforts for roots excavation, and decrease the time (and costs) of field occupation. It should be noted that the screening of field grown root systems at early stages is destructive, and that it can impede the final grain yield determination. However, this challenge can easily be circumvented by the inclusion of a few extra lines per plot during field establishment. Hence this enables the analysis of the RSA in a sub-plot, while keeping an undisturbed plot for later yield determination. The relatively short time needed to analyze one root system (Table 1) would make it possible to collect data from a large sample set, and time requirements would even further decrease when the manually determined traits (lateral density and secondary branching degree) could accurately be analyzed by additional imaging of a subsample or on few excised nodal roots, but such method remains to be developed and evaluated. Also when root excavation and washing could be mechanically assisted, throughput can be increased. The strengths, weaknesses, and future opportunities of this proposed method are presented in Table 6. Hereby it should be noted that, while this protocol is developed for rice, it could easily be transferred to other small-grained cereals that have a similar fragile root system, such as barley and wheat.

**Table 6:** Summary of the Strengths, Weaknesses, and Opportunities of this proposed phenotyping method for soil grown rice.

| Strengths | Weaknesses | Opportunities |
| --- | --- | --- |
| ➤ Robust protocol applicable at field scale<br>➤ Valid in remote environments<br>➤ Analysis of very complex root systems<br>➤ Cheap to develop and operate<br>➤ Fast data extraction<br>➤ Accessible to everyone<br>➤ (Semi-)Automated | ➤ Overlapping roots of very large plants<br>○ Can reduced accuracy of size-related traits<br>○ Short manual analysis needed<br>➤ Quantification of image-based traits remains challenging<br>➤ Destructive to the plant | ➤ Standardized protocol<br>○ Enables comparison among different experiments from different scientists<br>○ Contribution to a platform with standardized rice root data<br>➤ Faster analysis by mechanizing root excavation and washing<br>➤ Yield determination still possible in undisturbed sub-plot<br>➤ Determination of root angle possible on unwashed, excavated soil block<br>○ Method to be developed possibly by imaging technologies<br>➤ Easy transferrable to other small-grained cereals with fragile and dense root systems (e.g. wheat and barley). |

This suggested methodology is high-throughput, robust, low cost, and easy-to-learn, which indicates its large potential also in less-endowed environments. Future works would additionally benefit from developing a method that enables to determine the rice root angle on the excavated soil block before washing the soil (i.e. by scanning the position of the emerging nodal roots while rotating the excavated soil block), in order to complete the proposed protocol of rice root analysis in field conditions.

## 5 Conclusion

We evaluated two methods of root phenotyping for soil-grown rice (manual analysis versus image-based) and we propose an adjusted method, for inference on a minimum dataset to identify distinctive root phenotypes. After root excavating, the proposed method combines the manual determination of three traits that cannot be analyzed accurately by the imaging method only (i.e. the lateral density, the secondary branching degree, and the nodal thickness at the root base), with subsequent root imaging in a water bath followed by software-based image analysis (i.e. DIRT-software). This method should enable researchers to efficiently analyze the widest range of interesting root traits, achieve higher accuracy, reduce the required time of root system analysis, and it is applicable on fields in remote environments. This method would so enhance the collection of architectural root data of rice grown in agronomically relevant environments and the method enables the identification of different root phenotypes related to genotypes or induced by experimental treatments.

## 6 Declarations

### Ethics approval and consent to participate

Not applicable.

### Consent for publication

Not applicable.

### Competing interests

The authors declare that they have no competing interests. The authors also declare that the research was conducted in the absence of any commercial or financial relationships that could be construed as a potential conflict of interest.

## Author contributions

P.D.B., J.A., and R.V.H. conducted the experiments (with assistance as described in the acknowledgments), P.D.B. analyzed the data and wrote the manuscript. E.V., T.R., K.S., E.S., and R.M. conceived the research and revised the manuscript.

## Funding

This study was conducted at and partly financed by the Africa Rice Center in Tanzania and Madagascar and the KU Leuven in Belgium, and was additionally supported by the Flemish Interuniversity Council (VLIR-UOS) through a PhD scholarship (VLADOC grant) and a mobility scholarship. One experiment was financed by a C1 project (C16/15/042) funded by the KU Leuven.

## Acknowledgement

We thank Allen Lupembe and Leah Mwakasege for their assistance on the field and we thank all other Africa Rice staff who contributed to the work. We are grateful to Karlien Cassaert for the assistance in administration.

## Availability of data and materials

The datasets used and/or analyzed during the current studies are available from the corresponding author upon request.

## Notes

### Competing Interest Statement

The authors have declared no competing interest.

